# Abrupt light transitions in illuminance and CCT result in different temporal dynamics and interindividual variability for sensation, comfort and alertness

**DOI:** 10.1101/2020.11.19.389593

**Authors:** M.E. Kompier, K.C.H.J. Smolders, Y.A.W. de Kort

## Abstract

Detailed insights in both visual effects of light and effects beyond vision due to manipulations in illuminance and correlated color temperature (CCT) are needed to optimize study protocols as well as to design light scenarios for practical applications. This study investigated temporal dynamics and interindividual variability in subjective evaluations of sensation, comfort and mood as well as subjective and objective measures of alertness, arousal and thermoregulation following abrupt transitions in illuminance and CCT in a mild cold environment. The effects could be uniquely attributed to changes in illuminance or CCT and no interaction effects of illuminance and CCT were found for any of these markers. Responses to the abrupt transitions in illuminance and CCT always occurred immediately and exclusively amongst the subjective measures. Most of these responses diminished over time within the 45-minute light manipulation. In this period, no responses were found for objective measures of vigilance, arousal nor thermoregulation. Significant interindividual variability occurred only in the visual comfort evaluation in response to changes in the intensity of the light. The results indicate that the design of dynamic light scenarios aimed to enhance human alertness and vitality requires tailoring to the individual to create visually comfortable environments.

## Introduction

Light that is dynamic in illuminance and correlated color temperature (CCT) may induce ipRGC-influenced light responses (IIL; also known as non-image forming or non-visual effects) resulting in improved health, well-being and productivity of office workers (1). Several studies have investigated effects of such dynamic scenarios and hint towards positive effects on human functioning (e.g., 2–4). However, the lack of consolidated research methods prevents the formulation of a clear-cut conclusion (5). Preceding structured investigations of the overall effects of a daylong dynamic light scenario, the effects of the scenario’s individual elements (e.g., the transitions vs. the static parts) should be studied. In the current study, we aim to examine the temporal trajectories of alertness, arousal, comfort and mood – and individual differences therein – immediately following an abrupt transition in illuminance and/or CCT.

Effects of light on, for instance, sleep, human physiology, alertness and cognitive performance have been described for various exposure durations and levels of both illuminance (6–8) and CCT (9–11). Although the knowledge gained through these studies is very valuable, most studies do not provide detailed information about the short-term development over time of the various responses that a change in light induces (e.g., in terms of comfort, alertness, arousal and mood). Knowledge on the time it takes for an effect to emerge (**onset**) and whether the effect persists throughout the entire light exposure (**persistence**) is indispensable in the design of study protocols as well as practical light applications. This knowledge can be gained by continuous or repeated assessments of the dependent variable of interest after the onset of the light exposure. Although some laboratory studies did study participants’ responses to specific lighting conditions repeatedly on an hourly basis throughout the light exposure (12,13), more frequent repeated measures within one hour after onset of the light manipulation are also needed to determine these detailed temporal trajectories of the outcome parameters. Several studies reported that effects of a light manipulation on indicators of subjective alertness, physiological arousal and performance measures, if they emerged, were immediate and persistent (14–18). Others, however, reported delayed onset of performance effects (19,20) or self-reported alertness (21). One study reported significant effects of CCT on one of the performance tasks only mid-way throughout one hour of exposure (22). A last study (23) reported fast responses (within first 5 minutes) which then stabilized throughout 50 minutes for some measures (e.g., pupil response, EEG, heart rate), whereas other measures showed continued gradual incline or decline (e.g., HRV, DPG). The before-mentioned studies focused mainly on IIL responses and studied either illuminance or CCT. More knowledge on the response dynamics of both visual effects and effects beyond vision of combined manipulations of illuminance and CCT is needed to optimize study protocols as well as to design light scenarios for practical applications.

A recently published study examined response dynamics of visual experiences and light effects beyond vision–e.g., sensation, comfort, mood, self-reported alertness and physiology - after abrupt transitions in illuminance and CCT and reported markedly different trajectories for different indicators (18). Unfortunately, in this study changes in illuminances were always confounded with changes in CCT. Increases in both illuminance and CCT enhance the stimulation of the intrinsic photosensitive retinal ganglion cells (ipRGC) that are traditionally held responsible for the acute, alerting effects of light (24,25), but their effects on visual appraisals may work in opposite directions. Despite its alerting potential, cool, bright light is – at times – experienced as unsatisfying and uncomfortable (26), which then may also result in compromised alertness and performance (27). This demonstrates that the effects of illuminance and CCT may converge and hence strengthen each other for some measures, but diverge, potentially neutralizing each other for others. Studies investigating the effects of illuminance and CCT simultaneously yet independently are still quite scarce and generally do not focus on the temporal trajectories of the outcome variables in response to the manipulation of one or both of these parameters (26,28–31). This stresses the importance to study both light parameters simultaneously and disentangle these two factors.

To complicate matters, the literature on effects of illuminance and CCT has repeatedly suggested that the sensitivity for light responses is susceptible to individual differences. Although sometimes the lack or subtlety of effects of light on subjective sleepiness/alertness during daytime is attributed to interindividual differences (15,16,32), other studies have explicitly shown that substantial individual variability in responsiveness exists for different measures (e.g., visual comfort and circadian phase shifting responses of light) (33–35). Potential explanations for these individual differences in sensitivity to light could be the dependence on individual’s prior light exposure and thus be more situational (36,37), but it could also have a genetic basis and reflect trait rather than state differences (38,39).

In order to investigate the response dynamics of the independent and joint effects of illuminance and CCT, we studied the effects of abrupt step-wise increases in illuminance and CCT on participants’ evaluations of the ambient environment and their functioning. In this study we additionally examined the extent to which individual variability affects these responses. The employed study design also allows for a partial replication of the study by Kompier, Smolders, van Marken Lichterbelt and de Kort (18) on response dynamics after an abrupt transition in lighting. Despite the lack of effects on physiological and behavioral measures in Kompier et al. (18), these were included here to allow for replication of these null results. Whereas (18) was performed in thermoneutral conditions, the current study was performed in a mild cold environment as there are indications that slight discomfort may be a prerequisite for cross-modal interactions between thermal and visual comfort (40). The research question the current study sought to answer was how an abrupt transition from dim to bright light, or from warm to cool, influences subjective evaluations of comfort and mood as well as subjective and objective measures of alertness, arousal and thermoregulation over time in a mild cold environment. Moreover, we sought to investigate potential interactions between these interventions and study the nature and extent of interindividual variability in these effects.

## Materials and methods

### Design

Temporal responses to abrupt transitions in lighting were examined in a laboratory study with a two (Illuminance: bright vs. dim) by two (CCT: cool vs. warm) within-subjects design. Each 45-min condition was preceded by a 45-min baseline in warm, dim light (Fig 1) and presented during separate sessions, of which the order was partially counterbalanced. A combination of self-report, performance and physiological measures was used as outcome measures. The study was approved by the institutional ethical review board.

**Fig 1.**
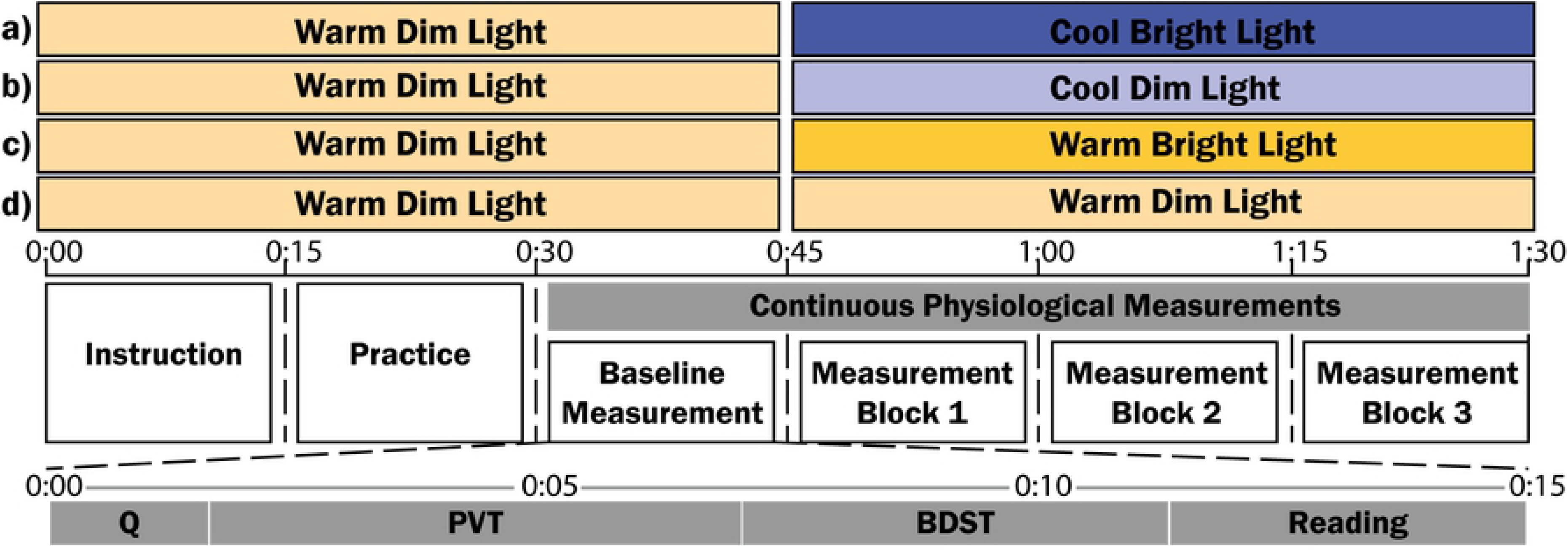
The four light conditions and the experimental procedure of one session. (A) cool, bright light (CB), (B) cool, dim light (CD), (C) warm, bright light (WB), and (D) warm, dim light (WD). Participants were exposed to these four conditions in separate sessions on different days in partial counterbalanced order. Q: Questionnaire, PVT: Psychomotor Vigilance Task, and BDST: Backward Digit Span Task

### Participants

Twenty-three healthy participants (13 female; M_age_ = 23, SD_age_ = 2.0; range = 18-26 years old) were recruited via the [REMOVED FOR BLIND REVIEW] database. None of the participants reported visual or auditory deficits, or were an extreme chronotype (assessed using the Munich Chronotype Questionnaire (41)). Additionally, none of the participants used medication other than the contraceptive pill or suffered from hypertension or cardiovascular disease. Last, none of them travelled intercontinentally in the past three months. Further descriptors can be found in Table 1. Participants gave their written informed consent and received a monetary compensation for their participation.

**Table 1.** Participant descriptors.

|  | Mean (SD) | *n* = 23<br>Range |
| --- | --- | --- |
| **Body Mass Index** | 22 (± 3.0) | 18 to 30 |
| **Midsleep (MSFsc)** | 4.7 (± 1.1) | 2.4 to 6.9 |
| **PSQI score** | 4.0 (± 2.4) | 1 to 10 |
| **General Health items (SF-36)** | 74 (± 14) | 45 to 100 |
| **Light Sensitivity – Eye Problems** | 1.8 (± 0.8) | 1 to 4 |
| **Light Sensitivity – Headache** | 1.7 (± 0.9) | 1 to 3 |
| **Light Sensitivity – Sunglasses** | 2.7 (± 1.3) | 1 to 5 |
| **Thermal Sensitivity** | 30 (± 1.1) | 1 to 4 |

### Setting and apparatus

An office setting with two separate desks was created in a climate chamber with a floor area of 18.0m^2^ and a ceiling height of 2.7m. With a Minolta Luminance Meter LS-100, the reflectance of the various surfaces in the room were measured (off-white walls: 80.9%, white partition wall: 91.8%, grey floor: 27.7%, light grey desk: 49.0%). The intended air temperature in the room was 17.0°C, but presumably due to the heat production of the participants the operative air temperature was 18.0°C ± .1 (air velocity = .01m/s ± .00; relative humidity = 67.4% ± 7.2, and black bulb temperature = 17.6°C ± .1). The four light settings were created using a set of four ceiling-mounted luminaires (PowerBalance Tunable Whites; RC464B LED80S/TWH PSD W60L60) above each desk. Table 2 shows the alpha-opic equivalent daylight illuminances (EDI), illuminance and CCT of all four light settings, and Fig 2 shows the spectral power distribution of the four light settings.

**Table 2.**
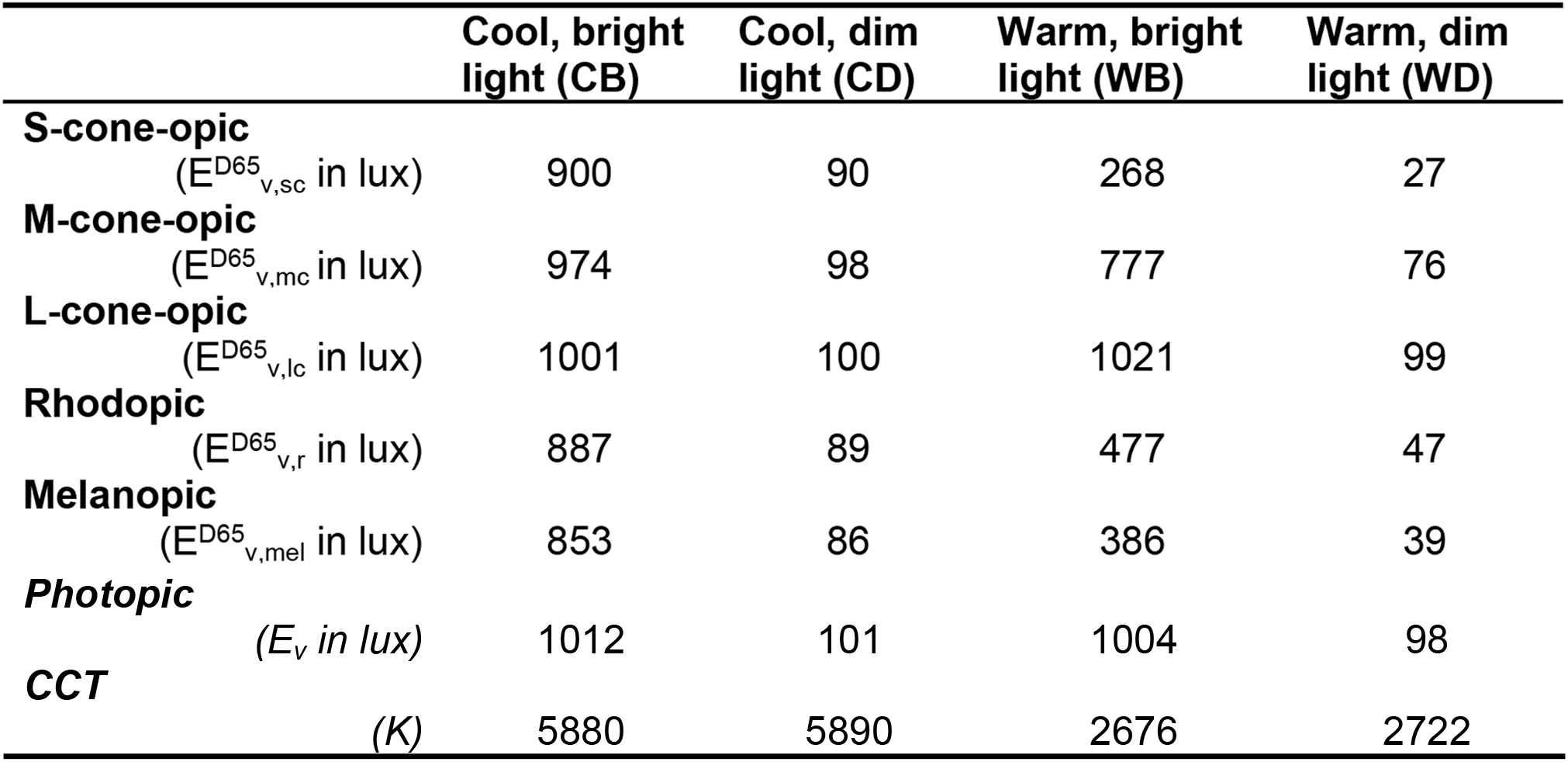
α-opic EDI, illuminance and CCT on the eye of the light settings.

**Fig 2.**
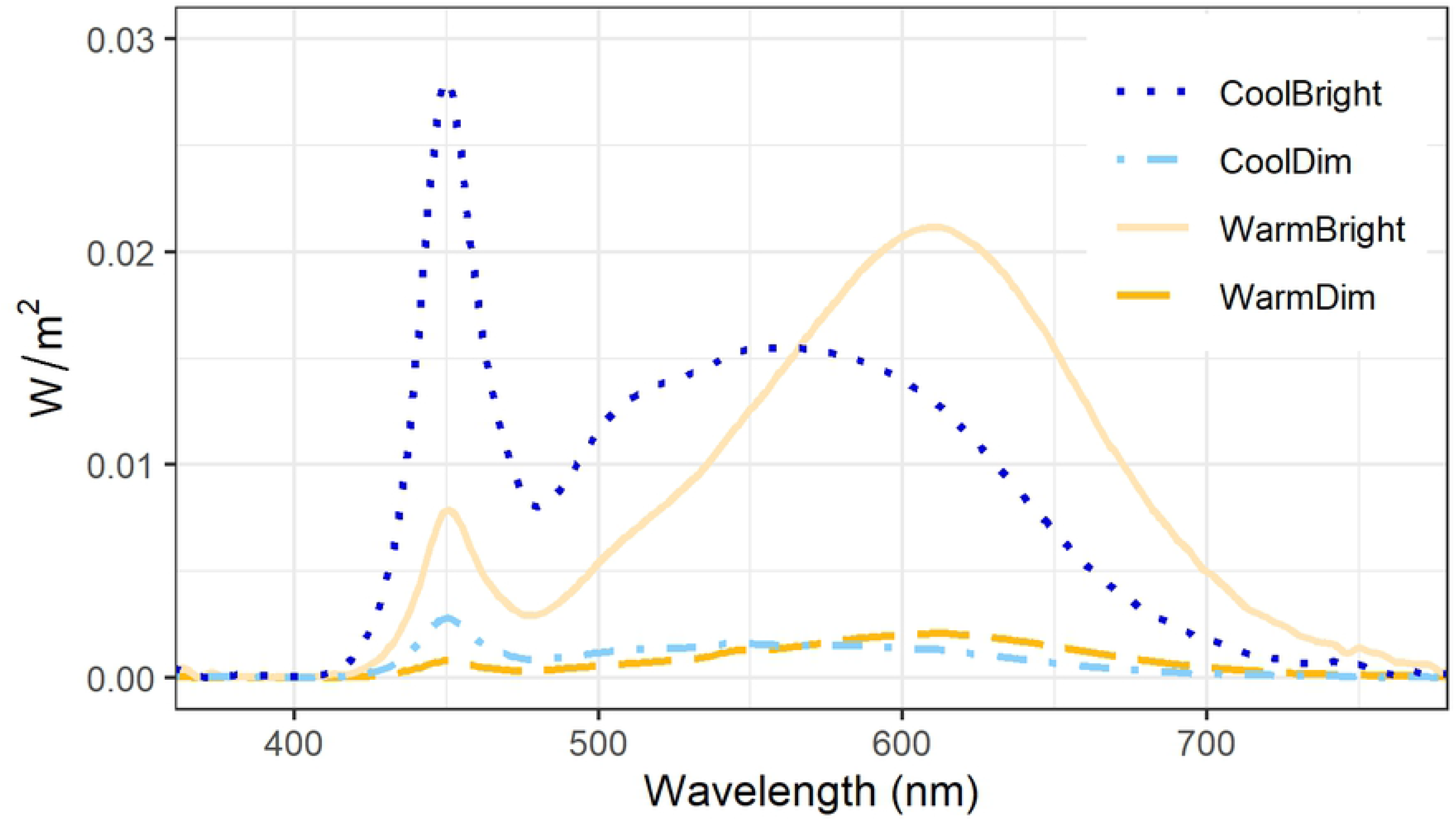
Spectral power distributions of the light settings.

### Measurements

#### Subjective measurements

Visual experience was probed as visual sensation, acceptance and comfort. Participants’ visual sensation was measured with two separate items for perceived brightness and color (Sensation_VI_ and Sensation_VC_) on 7-point rating scales ranging from *Very Low (−3)* to *Very High (3)* and *Very Cool (−3)* to *Very Warm (3)* respectively. Furthermore, participants evaluated acceptance of the visual environment (Acceptance_V_) on a binary scale (*Acceptable/Unacceptable*). Participants were also asked to assess comfort with the brightness and color of the white lighting respectively (Comfort_VI_ and Comfort_VC_) on 6-point rating scales ranging from *Very Uncomfortable (−2)* to *Just Uncomfortable (0)* and from *Just Comfortable (1)* to *Very Comfortable (3).* These two scores were averaged into Comfort_V_ as they had an internal consistency (Cronbach’s α) of .81.

Thermal experience was also probed with three items, which were based on ASHRAE Standard 55 (43). Thermal sensation (Sensation_T_) was evaluated on a 7-point scale ranging from *Cold (−3)* to *Hot (3)*, thermal acceptance (Acceptance_T_) on a binary scale (*Acceptable/Unacceptable*), and thermal comfort (Comfort_T_) on a 6-point rating scale ranging from *Very Uncomfortable (−2)* to *Just Uncomfortable (0)* and from *Just Comfortable (1)* to *Very Comfortable (3)*. Additionally, self-assessed shivering (SAS) was evaluated on a VAS ranging from *Not at all (1)* to *Yes, I shiver (10)*.

Subjective sleepiness was measured using the Karolinska Sleepiness Scale (KSS) with a response scale ranging from *Extremely alert (1)* to *Extremely sleepy (9)* (44). Vitality was measured using four items (Lively, Awake, Sleepy, and Drowsy; Cronbach’s α = .82) that were evaluated on 5-point scales ranging from *Definitely not (1)* to *Definitely (5)*. Vitality was calculated using the factor loadings that were derived through Principal Component Analysis (.70*Lively + .86*Awake - .82*Sleepy - .84*Drowsy). Participants evaluated their mood state (Tense, Calm, Sad, and Happy) on identical response scales. Tense and Sad were excluded from the analysis as the variance in the responses was too low. Calm was recoded into three categories (recode key: 1-3 = 1, 4 = 2 and 5 = 3) to correct for the skewed response distribution.

#### Performance tasks

In terms of performance we measured executive functioning and vigilance components. The total number of correct responses during the four-minute Backward Digit Span Task (BDST) was used as an indicator of executive functioning (45). Participants were presented with auditory sequences of numbers that ranged from four to eight digits. They completed two trials per digit-span length, resulting in ten trials per measurement block. After hearing the digits, participants had to type the sequence in reversed order within a time limit (2s + 2.3s per digit).

The mean reaction speed during the five-minute auditory Psychomotor Vigilance Task (PVT) was used as a measure of vigilance (46). Participants pressed the space bar of the keyboard as fast as possible in response to hearing short auditory stimuli of 400Hz with an inter-stimulus interval ranging between 6 and 25s.

#### Physiological measurements

Electrocardiography and electrodermal activity (EDA) measurements were done using TMSi software to collect physiological indicators of arousal. For heart rate (HR) and heart rate variability (HRV), participants attached one electrode on the left clavicula, one on the soft tissue below the right clavicula and one on the soft tissue right below the ribs on the left side of the body. The EDA electrodes were attached to the first phalanxes of the middle and ring finger of the non-dominant hand to measure skin conductance level (SCL) in micro Siemens. Institutional software was used for outlier and artefact detection (47–49). Mean HR, HRV (determined as root mean square of successive differences) and SCL during the five-minute PVT were used in the analyses.

Sixteen iButton (DS1925) dataloggers (sample interval 300s) attached using Fixomull Stretch tape were used to measure bodily temperatures. The iButtons were placed on 14 ISO-defined body sites (50), based on which mean skin temperature (T_average_) was calculated. Additionally, iButtons were placed at the under arm and the middle finger to assess peripheral vasoconstriction (51). The proximal skin temperature (T_proximal_) was computed by averaging the temperature measured at the scapula, paravertebral, upper chest, and abdomen. The distal skin temperature (T_distal_) was computed by averaging the fingertip, instep, hand, and forehead skin temperature. To avoid a disproportional distribution, fingertip and hand temperatures were averaged first. Mean skin temperature and the DPG (T_distal_ – T_proximate_) during the five-minute PVT were calculated to assess thermoregulation.

#### Control measurements

Light exposure on the day of the session was measured from awakening until the start of the session using a light sensor (LightLog). Furthermore, the Core Consensus Sleep Diary (42) was administered to assess sleep timing, duration and quality of the night before the experimental session. In addition, information on behavior, caffeine and food consumption was gathered at the start of the session (see S1). During the session, participants additionally evaluated the effort they had invested in the reading task and both performance tasks on a visual analogue scales (VAS) ranging from *None (0)* to *Very Much (20)*.

### Procedure

The study was conducted between May 24^th^ and June 24^th^, 2019. Participants completed all experimental sessions at the same time of day (8:45, 10:45, 13:30 or 15:30) to overcome time-of-day effects within participants. The sessions were generally scheduled at least one day apart to avoid session-to-session effects. Two participants had two of the four sessions on subsequent days due to practical limitations in the planning. Participants’ clothing was standardized at an estimated clothing value of .7 clo, including insulation of the chair.

After welcoming the participants, the adaptation period of 30 minutes in warm, dim light in the mild cold environment started. While adapting to this environment, control measures were taken and participants were instructed. Subsequently, participants applied the sensors for the physiological measurements and they practiced both the PVT and the BDST. After this adaptation period, the physiological measurements were started and continued for 60 minutes until the end of the session. During this period, the self-report measures and performance tasks were sampled every 15-min interval. The self-reports were completed in on average 84 ± 35s, after which the five-minute PVT and the four-minute BDST were performed. The first measurement, always in warm, dim light, was used as a baseline measurement. In the spare time throughout the procedure, participants read a book. The procedure, as shown in Fig 1, was identical for all sessions. At the end of the fourth session, participants were debriefed.

### Statistical analysis

MATLAB R2017b was used for all data processing and RStudio 1.1.463 for all analyses. The ‘psych’ and ‘plyr’ packages were used for the preparatory analyses; for the statistical analysis ‘emmeans’, ‘Hmisc’, ‘lme4’ and ‘lmerTest’ were used. The ‘ggplot2’-package was used for all visualizations. Outliers were identified by looking at the normal distributions and checking for cases that deviated more than three standard deviations from the mean. From the binary variables percentual scores were calculated and visual inspections were done instead of statistical analyses.

The main analyses examined the effect of *Illuminance* and *CCT* over time after the abrupt transition in lighting. In these mixed model analyses, Participant (P) and Session (S; nested within Participant) were added as random intercepts within the model. The models for each of the different dependent variables further included *CCT* (Warm/Cool), *Illuminance* (Bright/Dim), *Measurement block* (1/2/3), the two-way interactions *CCT*Illuminance*, *CCT*Block* and *Illuminance*Block*, the three-way interaction *CCT*Illuminance*Block* and *Time of day* (morning/afternoon) as fixed predictor variables. The *Baseline score* was added to account for potential baseline differences and *Reading effort* as an indicator of participants’ effort spent in between assessments. No other control variables were used as no statistically significant differences existed between neither the experimental conditions nor morning vs. afternoon sessions in the control variables assessed at session level. Furthermore, we examined whether random slopes for either Illuminance or CCT at the participant level significantly improved the model. Only in case the addition of the random slope resulted in a significant improvement compared to the model without random slope as determined by a likelihood-ratio test, the random slope was added to the final model. The model was specified as follows: *Y_ijk_ = β_0_ + P_00k_ + S_0jk_ + β_1_·Block + (β_2_+ P_1k_)·CCT + (β_3_+ P_2k_)·Illuminance + β_4_·CCT·Illuminance + β_5_·CCT·Block + β_6_·Illuminance·Block + β_7_·CCT·Illuminance·Block + β_8_·TimeOfDay + β_9_·Y_baseiine_ + β_10_·ReadingEffort + e_ijk_*. Based on the models, contrast analyses were performed to examine the onset and persistence of the effect of *Illuminance* and *CCT* separately per measurement block. For all analyses, an α of .01 was used as cut-off for statistical significance. In the results section, only the significant effects and differences are described for conciseness. Further statistics for the models without random slopes can be found in the S2-S4 Tables.

## Results

### Visual experience

Participants’ sensation of the brightness of the light (Fig 3A) was significantly affected by *Illuminance* (F(1,65) = 145.79, p<.001). As expected, the bright conditions were perceived significantly brighter (EMM ± SE = 1.36 ± .13) than the dim conditions (−.28 ± .13). Additionally, there was a significant main effect of *Measurement block* (F(2,182) = 13.46, p<.001) and an interaction of *Illuminance* and *Measurement block* (F(2,182) = 11.15, p<.001). Over the three measurement blocks, the brightness sensation of the bright conditions converged slightly towards the brightness sensation of the dim conditions. Yet, the effect of *Illuminance* emerged right after the transition (p<.001) and persisted throughout the remaining two measurement blocks (both p<.001). Participants’ sensation of the color of the white light (Fig 3B) was significantly affected by *CCT* (F(1,69) = 90.16, p<.001). In line with the expectations, the warm conditions were perceived as warmer (.29 ± .14) compared to the cool light conditions (−1.24 ± .14). In addition, there was a significant interaction effect of *CCT* and *Measurement block* (F(2,181) = 16.83, p<.001), reflecting a slightly diminishing difference between warm and cool conditions in the second and third measurement block compared to the first. Still, a statistically significant difference in sensation of the color of the light persisted across all measurement blocks (all p<.001).

**Fig 3.**
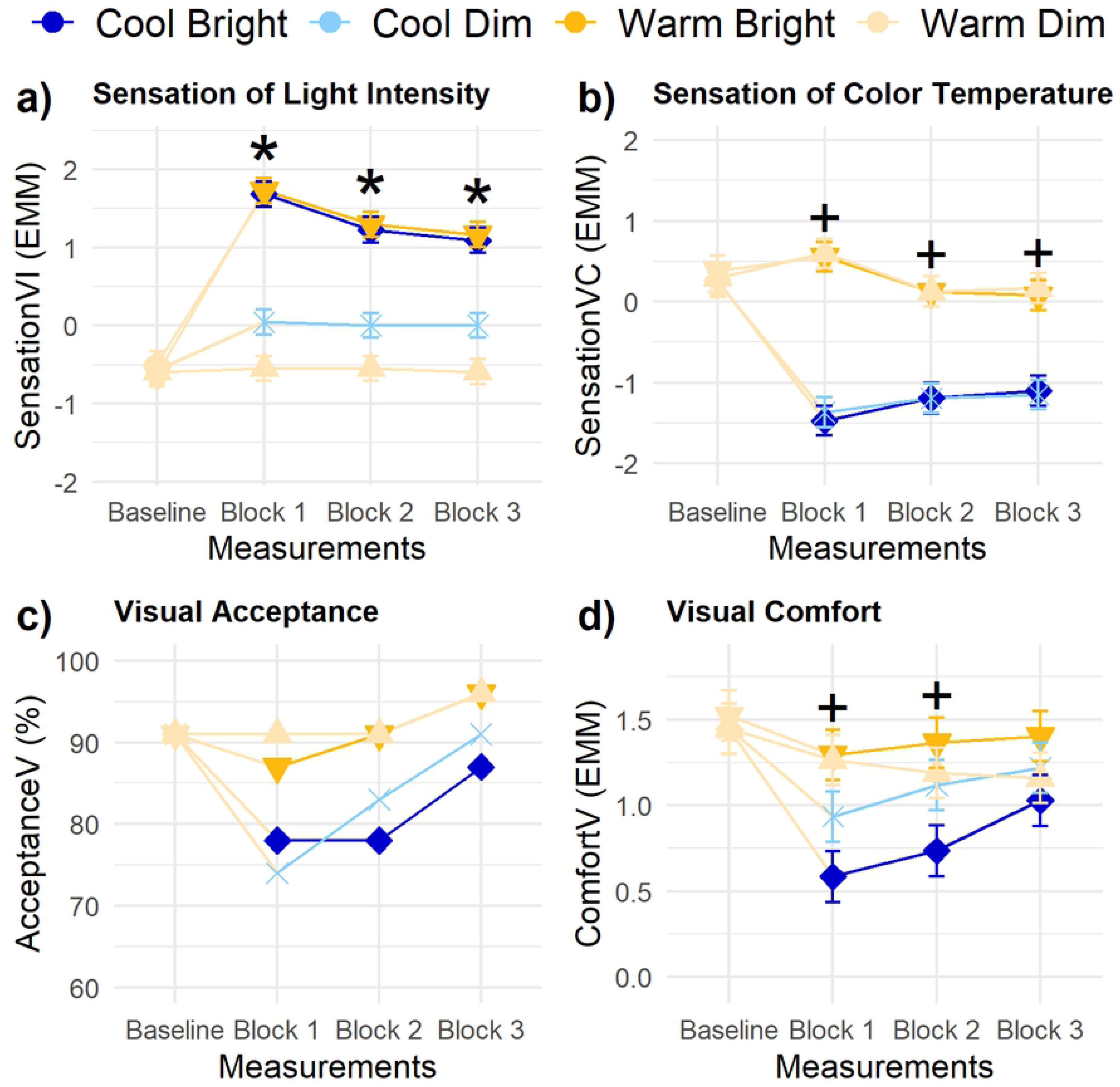
Trajectories of visual experience parameters. (A) sensation of light intensity, (B) sensation of color temperature, (C) visual acceptance (in % - no statistical testing), and (D) visual comfort. Error bars are standard errors (SE). Contrasts for illuminance and CCT were done for each measurement block: + indicates p<.01 for CCT, * p<.01 for illuminance.

Visual inspection of the results for visual acceptance suggest a lower visual acceptance of the light conditions after transitioning to the cool light conditions compared to both warm light conditions (Fig 3C). Over time, the percentage of persons rating the light setting as acceptable increased. For visual comfort, there was a statistically significant main effect of *CCT* (Fig 3D; F(1,44) = 13.42, p=.001), but not of *Illuminance*. However, the effect of *Illuminance* on Comfort_V_ was structurally influenced by interindividual differences (χ^2^ (2) = 18.82, p<.001); the illuminance of the light led to widely varying visual comfort votes by the participants (range = - .50 to 1.57). Fig 4A clearly shows that the bright light was evaluated as more comfortable than the dim light by some participants, but less comfortable by others. In contrast, the effect of *CCT* on ComfortV showed no statistically significant interindividual differences in responsiveness (χ^2^ (2) = .26, p=.88). Fig 4B indicates that the difference in CCT of the light conditions led to comparable changes in visual comfort evaluations for all participants; the warm light conditions were, on average, experienced as more comfortable (1.27 ± .10) compared to the cool light conditions (.92 ± .10). The effect of *CCT* on Comfort_V_ was only statistically significant in the first two measurement blocks after the transition (Block 1: p<.001, Block 2: p=.003, Block 3: p=.14). Last, *Reading effort* (β = -.03 ± .01, F(1,233) = 10.02, p=.002) and *Baseline visual comfort* (β = .28 ± .09, F(1,68) = 10.81, p=.002) had a significant effect on the visual comfort of the light condition. Comfort_V_ decreased with a higher self-reported reading effort.

**Fig 4.**
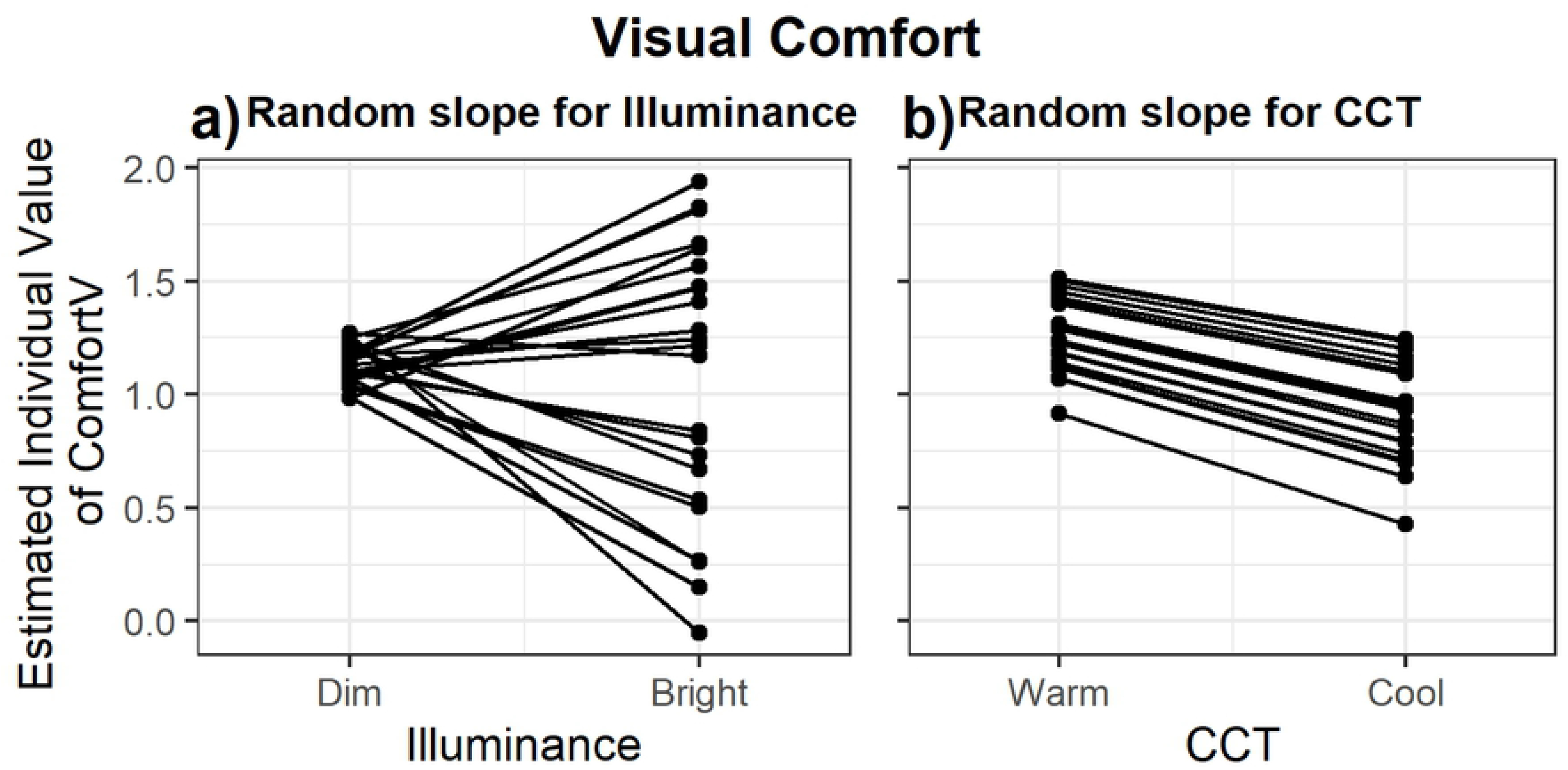
Individual differences in the visual comfort evaluation. Individual values of visual comfort are calculated using the estimated marginal mean + P_0j_ + P_1j_ * x based on the model with random slopes for (A) Illuminance and (B) CCT

### Vitality, sleepiness and performance

The significant main effect of *Illuminance* on vitality (Fig 5A; F(1,66) = 10.78, p=.002) indicated that vitality was, on average, higher in the bright light conditions (1.18 ± .29) compared to the dim light conditions (.18 ± .29). However, the contrast analyses revealed that bright light led to significantly increased vitality in the first two measurement blocks after the transition only (Block 1: p =.001, Block 2: p=.006, Block 3: p=.20). Random slope analyses showed no statistically significant interindividual differences. Vitality was significantly influenced by *Reading effort* (β = −.19 ± .03, F(1,226) = 44.50, p<.001) and *Baseline level of vitality* (β = .40 ± .07, F(1,90) = 30.16, p<.001). Vitality decreased with increasing self-reported reading effort. The response pattern for sleepiness was highly similar, but – as expected – reversed (Fig 5B). *Illuminance* had a statistically significant main effect on sleepiness (F(1,67) = 8.24, p=.005), indicating that the bright conditions were related to a lower sleepiness (4.57 ± .21) compared to the dim light conditions (5.22 ± .21). Again, this effect was only significant in the first two measurement blocks (Block 1: p=.004, Block 2: p=.006, Block 3: p=.23). Sleepiness was additionally affected by *Reading effort* (β = .11 ± .02, F(1,250) = 32.06, p<.001) and the level of sleepiness at baseline (β = .33 ± .08, F(1,97) = 15.66, p<.001). Sleepiness increased with a higher self-reported reading effort.

**Fig 5.**
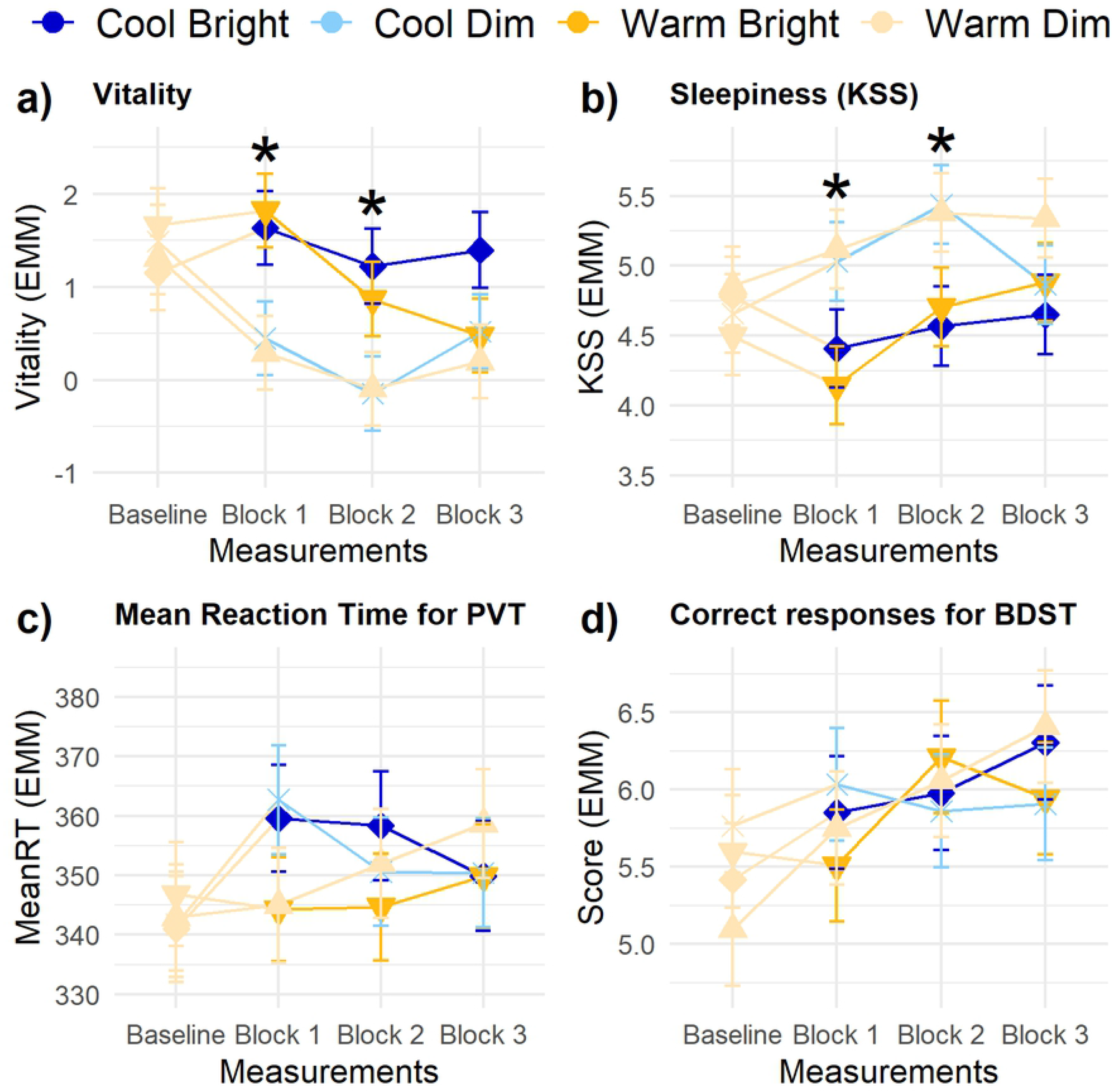
Trajectories of vitality, sleepiness and performance parameters. (A) vitality, (B) sleepiness, (C) mean reaction time for PVT, and (D) correct responses for BDST. Error bars are SE. Contrasts for illuminance and CCT were done for each measurement block: + indicates p<.01 for CCT, * p<.01 for illuminance.

The response time during the PVT showed neither statistically significant main nor interaction effects of *Illuminance* and *CCT* (Fig 5C), nor interindividual differences of these effects. *Baseline response time* significantly predicted the response time on the PVT (β = .51 ± .063, F(1,81) = 66.41, p<.001). Similarly, *Baseline BDST score* was a significant predictor for the number of correct responses on the BDST (Fig 5D; β = .35 ± .05, F(1,83) = 41.59, p<.001). No main or interaction effect of *Illuminance* and *CCT* on the score of the BDST occurred in any of the measurement blocks.

### Physiological arousal

SCL, HR and HRV were not significantly influenced by the light conditions (Figs 6A-C) nor by interindividual differences in the response to these light conditions. All three parameters were significantly affected by their *Baseline level* (β = 1.02 ± .05, F(1,69) = 465.50, p<.001; β = .89 ± .03, F(1,38) = 949.98, p<.001 and β = .87 ± .05, F(1,87) = 251.30, p<.001, respectively). SCL and HRV were, additionally, significantly influenced by *Measurement block* (F(2,164) = 18.48, p<.001 and F(2,169) = 28.76, p<.001, respectively), both showing an increase over time in the experimental session.

**Fig 6.**
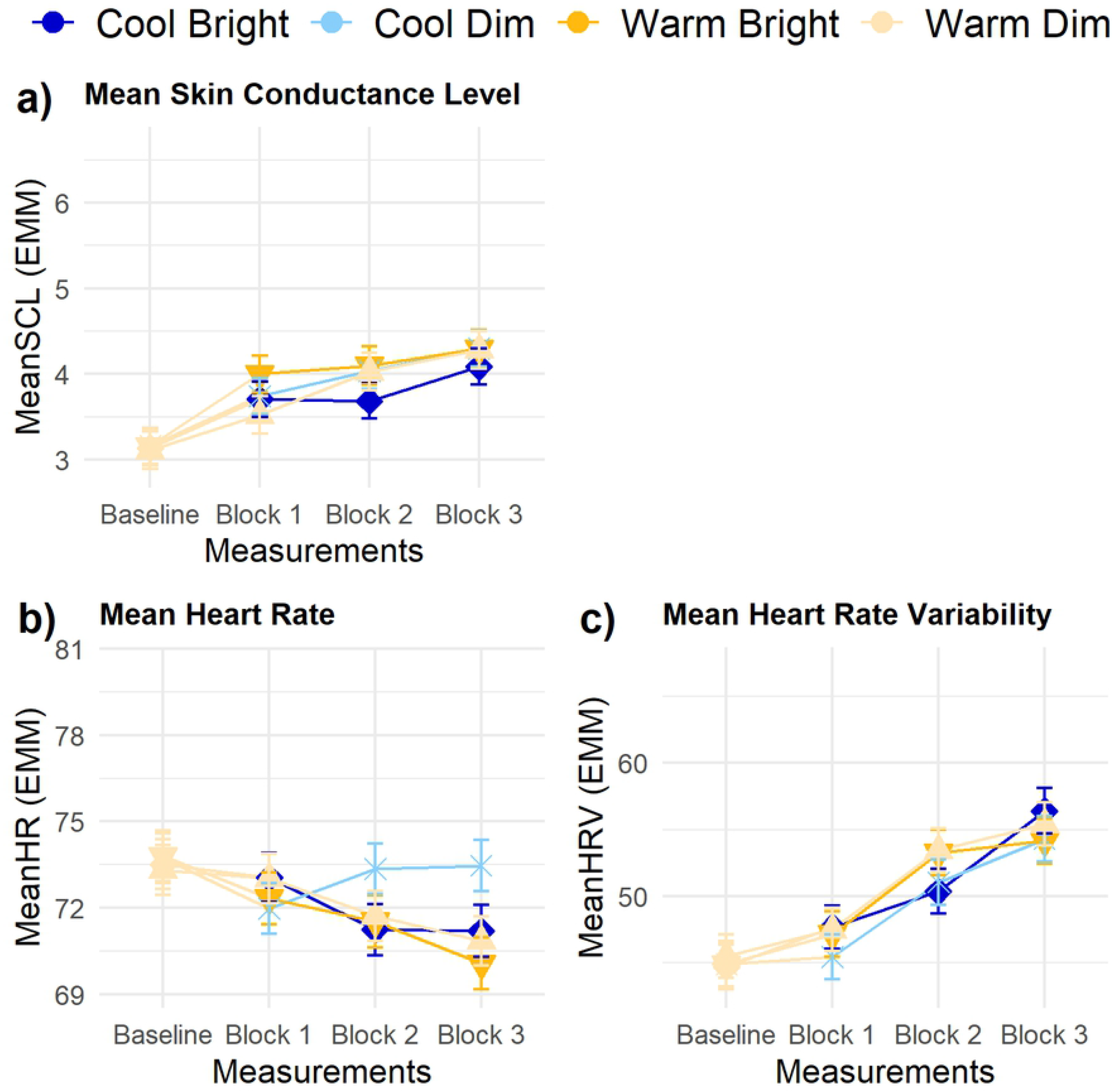
Trajectories of physiological parameters. (A) mean SCL, (B) mean HR, and (C) mean HRV. Error bars are SE. No significant differences existed for illuminance and CCT in any of the measurement blocks.

### Mood

Participants’ calmness ratings were significantly influenced by *Baseline level* (β = .50 ± .07, F(1,94) = 45.26, p<.001) and the three-way interaction of *Illuminance, CCT* and *Measurement block* (F(2,183) = 4.94, p=.008) suggesting slightly different trajectories for all conditions (Fig 7A). Yet, no main and interaction effects of *Illuminance* and *CCT* in these responses occurred in any of the measurement blocks. Happiness showed an significant effect of its baseline score (Fig 7B; β = .61 ± .08, F(1,87) = 54.58, p<.001). In the first measurement block, there was also a significant difference between the warm and cool conditions: the transition to cool light led to a decrease in reported happiness (p=.007) in the first measurement block. These effects turned insignificant in the following two measurement blocks (Block 2: p=.25, Block 3: p=.56).

**Fig 7.**
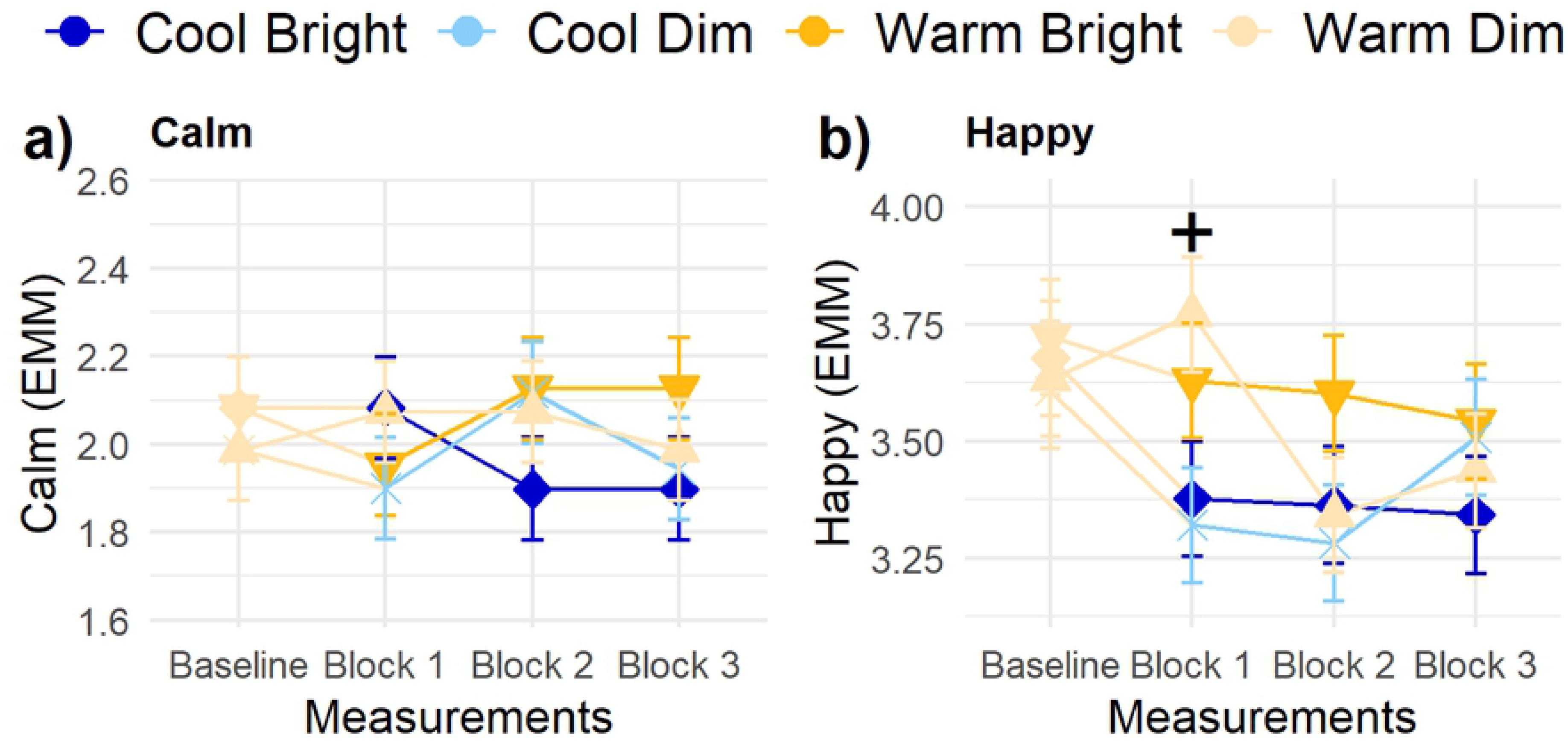
Trajectories of mood parameters. (A) Calm, and (B) Happy. Error bars are SE. Contrasts for illuminance and CCT were done for each measurement block: + indicates p<.01 for CCT, * p<.01 for illuminance.

### Thermal experience

Thermal sensation of the participants (Fig 8A) was significantly influenced by *Measurement block* (F(2,182) = 11.76, p<.001) and *Baseline thermal sensation* (β = .35 ± .09, F(1,87) = 16.60, p<.001). Sensation_T_ decreased over time, but in none of the measurement blocks a significant main or interaction effect of *Illuminance* or *CCT* occurred (all p > .01). Similarly, self-assessed shivering (Fig 8B) was only significantly affected by *Measurement block* (F(2,177) = 9.16, p<.001) and *Baseline level* (β = .46 ± .13, F(1,75) = 13.47, p<.001). Self-assessed shivering increased over time in session. Visual inspection of the percentage of persons rating the lighting conditions as thermally acceptable (Fig 8C) suggested a decrease over time in all conditions. The visual inspection suggests a lower thermal acceptance in the warm light conditions, however, this difference already seemed to exist in the baseline. Comfort_T_ (Fig 8D) was significantly affected by *Baseline level* (β = .21 ± .08, F(1,84) = 6.61, p=.01) and *Reading effort* (β = −.04 ± .01, F(1,266) = 10.33, p=.001). Participants’ thermal comfort significantly decreased over time and with increasing reading effort. The thermal experiences were not affected by the changes in the light conditions and showed no statistically significant interindividual variability in their responses to the lighting conditions.

**Fig 8.**
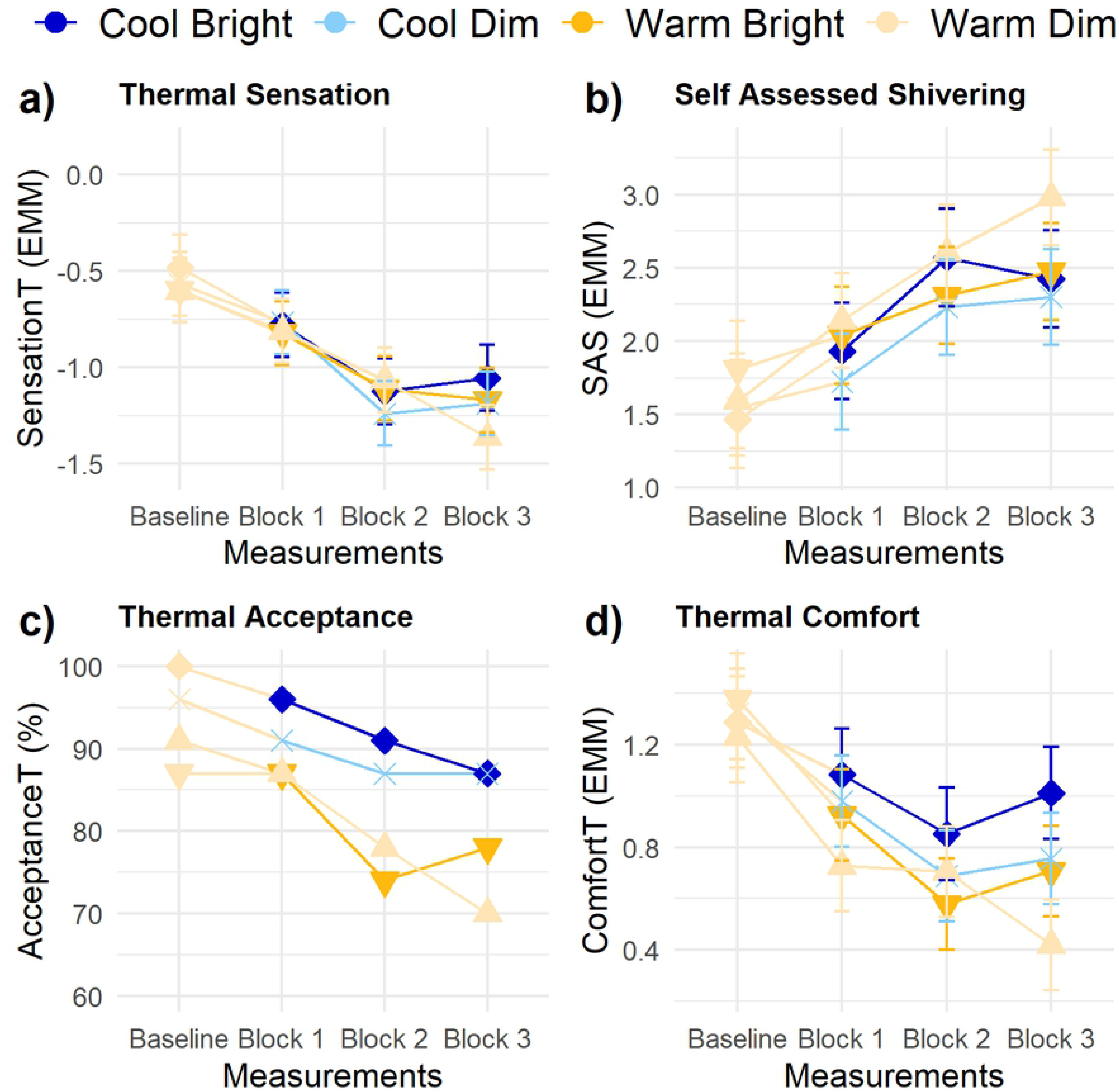
Trajectories of thermal experience parameters. (A) thermal sensation, (B) self-assessed shivering, (C) thermal acceptance (in % - no statistical testing), and (D) thermal comfort. Error bars are SE. No significant differences existed for illuminance and CCT in any of the measurement blocks.

### Thermoregulation

For the average skin temperature, there was no statistically significant main or interaction effect of *Illuminance* and *CCT* in any of the measurement blocks (Fig 9A), nor were there statistically significant interindividual differences in these effects. The average skin temperature was significantly affected by *Measurement block* (F(2,182) = 298.73, p<.001) and *Baseline skin temperature* (β = .98 ± .03, F(1,93) = 805.83, p<.001). Average skin temperature decreased over time. Similarly, the DPG (Fig 9B) was significantly influenced by *Measurement block* (F(2,183) = 22.55, p<.001) and *Baseline DPG* (β = 1.14 ± .03, F(1,88) = 1852, p<.001). Over the three measurement blocks, the DPG decreased (i.e., the difference between distal and proximal temperatures increased).

**Fig 9.**
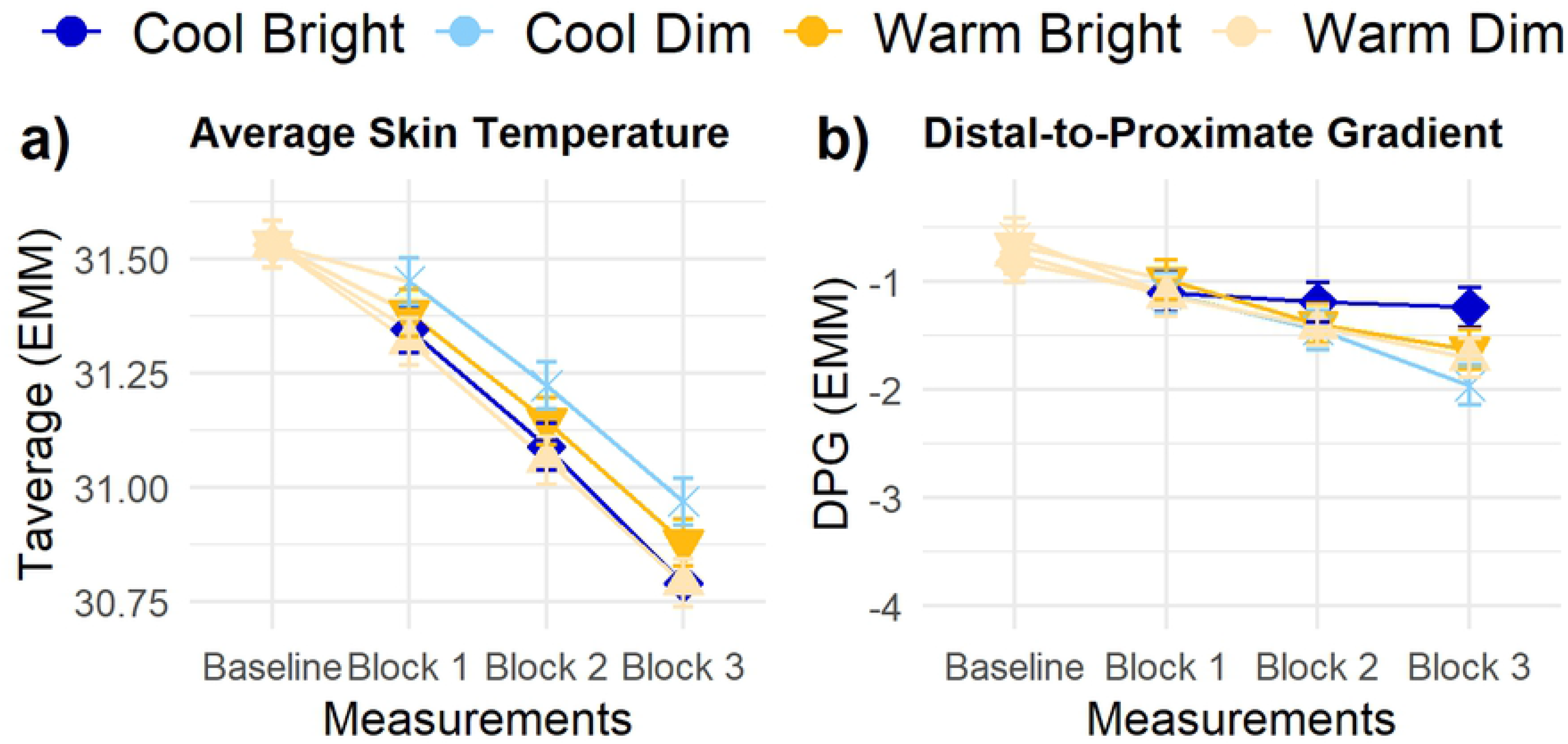
Trajectories of thermoregulation related parameters. (A) average skin temperature, and (B) distal-to-proximate gradient. Error bars are SE. No significant differences existed for illuminance and CCT in any of the measurement blocks.

## Discussion

With this laboratory study, we aimed to study the impact of an abrupt transition in illuminance and CCT on temporal trajectories of alertness, arousal, comfort and mood. Furthermore, we investigated the presence of interindividual variability in these response patterns. Healthy participants were exposed to four conditions and repeatedly completed subjective measures of comfort, alertness and mood, and performance measures probing vigilance and executive functioning, while continuously tracking physiological indicators of arousal and thermoregulation. We largely replicated findings reported by Kompier et al. (18), but – due to the adapted design of the current study – effects can now be uniquely attributed to changes in illuminance or CCT. The results provide insights on how people respond to abrupt transitions in illuminance and CCT and can be implemented in the design of daytime dynamic light scenarios.

### Onset - acute effects of an abrupt transition

The abrupt transitions in illuminance and CCT led to an immediate change in brightness and color sensation. Whereas the change in brightness perception could be clearly attributed to change in illuminance of the light, both color sensation and visual comfort showed a main effect of the transition in CCT only. More specifically, participants reported diminished visual comfort directly after the transition to cool light compared to the warm light conditions. Furthermore, participants reported feeling less happy right after transitioning to cool light compared to warm light. Significant decreases in sleepiness and increases in vitality also occurred immediately after the transition to bright light compared to the continuous dim light, which is in line with prior research (18,20). In contrast, none of the task performance and physiological arousal parameters showed either a direct or delayed response within 45 minutes after the abrupt transition in illuminance and/or CCT. A potential reason for the lack of a response to the light conditions in terms of task performance could be the compensation for the effect by the exerted effort on the task (52). Moreover, the absence of effects of different light conditions on performance and physiological arousal measures is not uncommon, and prior studies have reported that effects on these measures may depend on certain times of day, exposure duration, task difficulty, or season (12,15,19,53). Although this could explain why light did not affect these responses in the present study, we should question whether the acute effects of light on performance tasks and physiological arousal are practically relevant if they only occur in specific contexts. Last, in this study no direct evidence for cross-sensory effects of illuminance or CCT on thermal experience was found, although various theories (e.g., hue-heat hypothesis (54), brightness associations (55) or physiological effects (56)) hypothesize such effects. In sum, this study demonstrated that subjective responses to an abrupt transition in illuminance and/or CCT, if they emerge, always emerge virtually immediately and that task performance and physiological measures do not respond within the first 45 minutes after an abrupt transition.

### Persistence – sustained effects of an abrupt transition

Not all effects that occurred right after the transition persisted throughout the entire duration of exposure. Only the sensation of the color and brightness of the light remained significantly different between conditions across all three measurement blocks, even though the difference diminished slightly. As in Kompier et al. (18), the contrast between the persistent response for the visual sensation and the transient response for visual comfort and acceptance can be explained by adaptation to the light setting. Although participants’ sensation of the light persisted, the adaptation of the eye likely led to altered acceptance and comfort votes. The abrupt transition to cool light led to an immediate decrease in visual comfort, but this effect disappeared over time. This is partially in contrast to the review by Fotios et al. (57) who discussed that pleasantness ratings may reach a plateau at 6 minutes after light onset in certain settings, but also demonstrated that overall the effects of CCT on ratings of brightness and pleasantness are small and seem largely independent of adaptation time. After the light transition to cool, bright light, the visual comfort of the participants improved gradually. This suggests that initially participants may not appreciate an abrupt transition to cool light, but when time passes their visual comfort seems to returns towards the original level. Potentially, more gradual light conditions can prevent this initial transient decline in comfort. Happiness also responded temporarily to the transition in CCT, which was not in line with Kompier et al. (18), who found no effects of a transition from warm, dim to cool, bright lighting on happiness. The effects of the light conditions on vitality and alertness, on the other hand, were in line with earlier studies (14–18). The effects of illuminance on vitality and alertness persisted, at least throughout the first 20 minutes, after the transition. In this study, we demonstrated that the change in illuminance of the light, rather than the change in CCT, was significantly related to a change in sleepiness and vitality. The *E^D65^_v,mel_* of the bright compared to the dim conditions implied a tenfold increase, whereas the increase ‘merely’ doubled from the warm to the cool condition. In line with this, the – nonsignificant – difference in sleepiness and vitality between warm vs. cool light was about one fifth of the size of the effect of illuminance. Rather than attributing the alerting effect to illuminance or CCT per se, the melanopic activation that a light setting can achieve is the more likely explanation for this acute, alerting effect (58,59), as was also suggested in the study by Ru et al. (31). In this respect, it is important to note that the factor two difference in *E^D65^_v,mel_* between the warm, bright and the cool, bright light condition did not lead to a difference in sleepiness or vitality, which is possibly due to the nonlinear (sigmoidal) relationship between light and alerting responses (59,60).

### Interindividual variability

In this study, we additionally examined whether significant interindividual differences exist for the main effects of illuminance and CCT. To test the existence of this variability, random slopes for illuminance and CCT were added separately at the participant level in the models. The interindividual differences reported here therefore reflect variation between persons that is stable throughout the period during which participants completed the four sessions. More specifically, this variability is likely trait-like and potentially a function of person characteristics, such as lens characteristics or chronotype. Additionally, variation in responsiveness light may also emerge due to behavioral and situational differences, such as prior light exposure (37), but these were not examined in the current study. Interindividual variability was found, and exclusively so, for the effect of illuminance on visual comfort, which might explain the absence of a main effect of illuminance on visual comfort. Both the direction and the extent to which individuals’ visual comfort was affected were highly dependent on the individual; the transition to a higher illuminance was considered more comfortable by some and less comfortable by others. Examining the origin of this interindividual variation in responses to light requires between-subject comparisons which were not performed in this study due to statistical power considerations. Although individual variability in preferred lighting conditions have been demonstrated before (33,61), the current study is – to our knowledge – the first to demonstrate that individual differences for the effect of illuminance on visual comfort exist, but not for the effect of CCT. This finding, of course, applies only to the employed target audience, stimulus range, duration and timing, and requires further investigations in order to generalize. No significant interindividual differences in the alerting responses were found, despite suggestions thereof in prior studies (16,32). This might be explained by the homogeneity in the sample; limited variation in factors such as age and chronotype likely results in little variation in responsiveness to light (62).

### Theoretical reflections and practical implications

In the current study, we successfully disentangled the effects of illuminance and CCT on measures of subjective experiences and comfort. The results indicated that manipulations of illuminance and CCT did not interact in their alerting effects, and effects could be attributed to illuminance only. However, the difference in melanopic activation due to the contrasting light conditions may very well be the driving force behind the significant effect that was found (59). Although effects of illuminance and CCT on visual sensation were straightforward and consistent between persons, visual comfort showed marked individual differences. This means that increases in illuminance and CCT may result in a decline in visual comfort for some people, but have contrasting effects for others. None of the other outcome parameters that were included in this study showed interindividual variability in their responses to changes in illuminance or CCT, indicating similar responses to light for the relatively homogenous sample that was included in this study. Furthermore, the absence of effects on performance tasks, physiological arousal and thermal experience indicated that no overall daytime effects of light on these parameters emerged, at least not within 45 minutes of exposure, which is largely consistent with recent reviews (39,63).

The different parameters that relate to alertness, arousal and vigilance showed varying response patterns, demonstrating the nuanced differences between how subjective alertness, physiological arousal and behavioral vigilance can be influenced. This emphasizes that interpretation and explanation of effects of light on these parameters and their underlying mechanisms requires a multi-measure approach (64). Last, the occurrence of both transient and persistent responses underlines the delicacy in timing of measurements within a measurement protocol.

### Limitations and future research

Although this study was conducted in a relatively controlled environment, some limitations could not be overcome. In the climate chamber, two office-like workplaces were created to assure that two participants could participate simultaneously. Despite the use of a partitioning wall, participants may have been aware of each other, which potentially impacted concentration on tasks. Furthermore, the sample was predominantly young and healthy, which limits generalizability. The homogeneous sample may have also resulted in the inability to identify interindividual variability for some responses as these may depend on factors such as chronotype and age which showed little variation in our participant sample. To be able to generalize the results to the general population, larger and more heterogenous samples should also be included in future studies. Participants completed all sessions at the same time of day to control for variation due to time of day within participants, but interaction effects due to time of day between participants could not be tested due to power considerations. Future studies should investigate the effect of seasonality and time of day on the results (15,53,65) and examine the current effects in more naturalistic settings using longer exposure durations before the results can be applied in a dynamic light scenario for office environments. Last, we suggest that future research investigates more closely how visual comfort evolves over the day, extending the work by Kent, Altomonte, Wilson and Tregenza (66) on discomfort glare to visual comfort, to see whether dynamically changing light conditions throughout the day could be stimulating and lead to increased visual comfort.

### Conclusions

In the current study, we demonstrated that abrupt transitions in illuminance and/or CCT led to different response dynamics for subjectively evaluated appraisal, mood and alertness. Illuminance and CCT did not structurally interact on these markers, but each affected a selection (e.g., vitality and sleepiness were moderated by illuminance and visual comfort by CCT). Responses to the abrupt transitions in illuminance and CCT occurred always immediately and exclusively amongst the subjective markers. No obvious responses were found in objective measures of vigilance or arousal. Although subjective alertness benefitted immediately and for at least 20 minutes from higher light levels, we saw no measurable effect for physiological arousal and task performance. Thermal comfort and thermoregulation were also not significantly influenced by the lighting manipulations. In visual comfort evaluations (and only there) we detected statistically significant interindividual differences in responses to changes in the intensity of the light. The results indicate that the design of dynamic light scenarios that are beneficial for human alertness and vitality requires tailoring to the individual to create visually comfortable environments.

## Acknowledgements

We would like to thank Kay Tuip for his help in gathering the data, the HTI lab support team for the technical assistance of the materials and Wout van Bommel for the technical assistance in the laboratory.

## Supporting information

**S1 File. Start questionnaire.** The questions posed at the start of the session as control measures.

**S2 Table. Full model statistics.** Variances, model fit and R^2^ of the random intercept models.

**S3 Table. Estimated marginal means for main effects.** Delta, t-ratio and p-value for the main effect of illuminance and CCT separately.

**S4 Table. Estimated marginal means per measurement block.** Delta, t-ratio and p-value per measurement block for the effect of illuminance and CCT separately.

